# Sex differences in deleterious genetic variants in a haplodiploid social insect

**DOI:** 10.1101/2021.08.26.457852

**Authors:** Sara E. Miller, Michael J. Sheehan

## Abstract

Deleterious variants are selected against but can linger in populations at low frequencies for long periods of time, decreasing fitness and contributing to disease burden in humans and other species. Deleterious variants occur at low frequency but distinguishing deleterious variants from low frequency neutral variation is challenging based on population genetics data. As a result, we have little sense of the number and identity of deleterious variants in wild populations. For haplodiploid species, it has been hypothesized that deleterious alleles will be directly exposed to selection in haploid males, but selection can be masked in diploid females due to partial or complete dominance, resulting in more efficient purging of deleterious mutations in males. Therefore, comparisons of the differences between haploid and diploid genomes from the same population may be a useful method for inferring rare deleterious variants. This study provides the first formal test of this hypothesis. Using wild populations of Northern paper wasps (*Polistes fuscatus*), we find that males have fewer overall variants, and specifically fewer missense and nonsense variants, than females from the same population. Allele frequency differences are especially pronounced for rare missense and nonsense mutations and these differences lead to a lower genetic load in males than females. Based on these data we estimate that a large number of highly deleterious mutations are segregating in the paper wasp population. Stronger selection against deleterious alleles in haploid males may have implications for adaptation in other haplodiploid insects and provides evidence that wild populations harbor abundant deleterious variants.

## Introduction

Mutations are the raw source of variation for populations and most non-neutral mutations are deleterious (1). Identifying deleterious and disease-causing variants and distinguishing these variants from neutral genetic variation is of major interest for understanding disease and for predicting the evolutionary response of populations (2–4). Most deleterious variants are derived alleles present at low frequencies within populations, however new neutral mutations share these same characteristics (5–7). As a result, inferring the fitness effects of rare genetic variants from population genomics data remains a challenge, especially for non-model organisms (7, 8).

In arrhenotokous haplodiploid species, males develop from unfertilized eggs as haploids whereas females develop from fertilized diploid eggs. In essence, the entire haplodiploid genome is comparable to sex chromosomes in diploid species (9). This naturally occurring variation in ploidy between haplodiploid sexes has led to a long-standing yet untested hypothesis that deleterious alleles will be directly exposed to selection in males, but in females deleterious alleles can be masked from selection by partial or complete dominance (10–12) (Figure 1). Thus, in haplodiploid species, selection against deleterious alleles is predicted to occur more frequently in males than females, and is expected to lead to greater male mortality as a consequence of the purging of lethal recessive mutations (13). Identifying which genetic variants are differentially removed in haplodiploid males should therefore give insight into the evolutionary dynamics of deleterious mutations.

**Figure 1:**
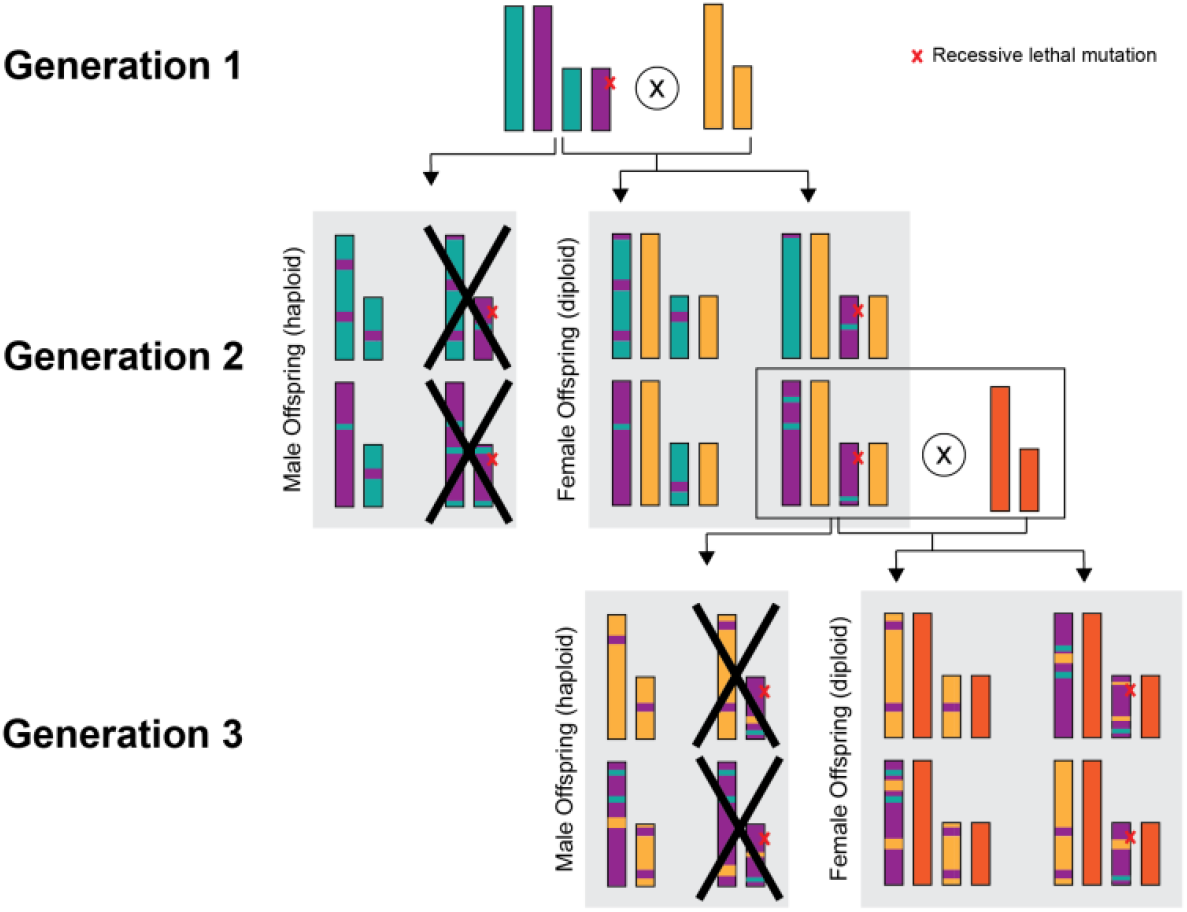
Schematic of differential selection against a recessive lethal mutation in an arrhenotokous haplodiploid species (N=2). All males with the recessive lethal mutation die but females must have two copies of the allele to be affected. After three generations, fifty percent of male offspring have been removed from the male gene pool while there was no selection against female offspring with the allele. No recombination occurs in haploid males.

In this study, we leveraged whole genome re-sequence data to directly test the hypothesis that deleterious alleles are preferentially purged in haplodiploid males. There are several predictions following from this hypothesis. First, samples of males should have fewer variants on average than samples of females from the same population. However, a natural consequence of haplodiploidy is that at equal sex ratios, females have a greater effective population size because females have double the number of copies of each chromosome (12), and consequently, females are expected to have twice the effective mutation rate as males. Therefore, differences in the number of variants between sexes could also result from an increased input of new mutations in females, rather than the removal of deleterious alleles in males. Mutations are random throughout the genome; therefore, a second prediction of this hypothesis is that if allele frequency differences between sexes are driven by selection, deleterious variants should be underrepresented in males relative to females. Deleterious variants are those which reduce the fitness of an organism. Across organisms, sites that modify protein amino acid sequences or influence transcription are more conserved, indicating they are under stronger purifying selection. Therefore, we specifically expect fewer nonsense or missense variants in males compared to females as these are most likely to have negative fitness consequences. Finally, if deleterious variants are more often removed from the male population, then strongly deleterious alleles will be absent or at a very low frequency in males but may be present at a higher frequency in the female population. As a result, we would expect an overall difference in allele frequencies between the sexes, and this difference should be most pronounced for deleterious variants as neutral low frequency variants are predicted to have a similar frequency in both males and females.

While there have been no direct tests of this hypothesis in haplodiploids, several lines of indirect evidence support differences in selection rates between haploids and diploids. In plants, purifying selection was greater on genes expressed only during the haploid phase than on genes expressed only during the diploid phase (14). Haplodiploid insects have lower genetic variability (15) and reduced inbreeding depression than diploid insects (16), consistent with haplodiploid species being more efficient at purging deleterious mutations and thereby having a lower genetic load relative to diploid species (17). Lastly, within haplodiploid species, if deleterious alleles are expressed more often in males, we might predict that on average, males will be less capable of coping with agents of selection. Accordingly, haplodiploid insect males have lower tolerance to pesticides than females (18), and male parasitoid wasps have been shown to have higher mortality rates than females (13).

We directly investigated this hypothesis using the Northern paper wasp, *Polistes fuscatus*, a primitively eusocial arrhenotokous haplodiploid wasp with a widespread distribution across the Eastern United States. In this species, reproductive males and reproductive females (gynes) are produced at an approximately 1:1 ratio in the fall (19). Males disperse from their natal nests to nearby lekking sites to mate, dying prior to winter. Gynes overwinter in hibernacula and start new nests the following spring (20, 21).

We collected 100 female and 185 male *P. fuscatus* from sites around Ithaca, New York from 2015-2019. To account for possible differences in the per chromosome *de novo* mutation rate between sexes, our dataset roughly balanced the number of chromosomes from males and females rather than the number of individuals. Wasps from this region form a large single panmictic population with a high degree of gene flow among sampling sites (22). All males and most females were reproductive individuals collected in the fall (Table S1). Within each sex, individuals were unrelated, however males and females from the same nest were included in the dataset. This heterogeneity in collection year and sampling site within our dataset should, if anything, make it more challenging to detect systematic allele frequency differences between the sexes.

Individually barcoded paired-end whole genome libraries were sequenced and aligned to the *P. fuscatus* reference genome (23). The dataset was filtered to remove variants with low depth of coverage to eliminate non-biological variants caused by sequencing errors as well as variants with unusually high coverage to remove variants from unresolved duplicated regions caused by genome assembly errors. An unavoidable consequence of haplodiploidy is that it is easier to detect true variants in males than females. At the same sequencing depth, sequence data from females is split between the two copies of each chromosome, while sequence data from males is from a single chromosome. Furthermore, heterozygous genotypes in males are caused by sequencing errors, decreasing the rate of false positives in males. Whereas females have a higher false negative rate due to an increased difficulty in distinguishing between sequencing errors and heterozygote genotypes, particularly at lower coverage sites. This difference in variant detection error rates between males and females is expected to increase the number of high confident variants in males relative to females, biasing the dataset towards a greater number of male variants, which is in the opposite direction predicted by our hypothesis.

### Females have more variants than males

We identified 4.6 million SNPs in the female samples and 3.6 million SNPs in male samples, matching our prediction that females will have more variants than males. However, due to the heterogeneity in sample size and sampling regime between the sexes, it may be more meaningful to compare the relative number of variants between the sexes rather than the total number of variants to pinpoint which regions of the genome have stronger purifying selection in males. We calculated nucleotide diversity (π) and SNP density for males and females, plotted these values in 50,000 bp windows across the genome, and calculated a z-score using the residuals between each point and the regression line (Figure 2). Using a cutoff of z>3, there were 74 outlier windows (1.7% of windows) for π, and 30 outlier windows (0.8% of windows) for SNP density. In all π outlier windows, females had higher than expected π. Similarly, 90% of SNP density outlier windows had greater SNP density in females than males.

**Figure 2:**
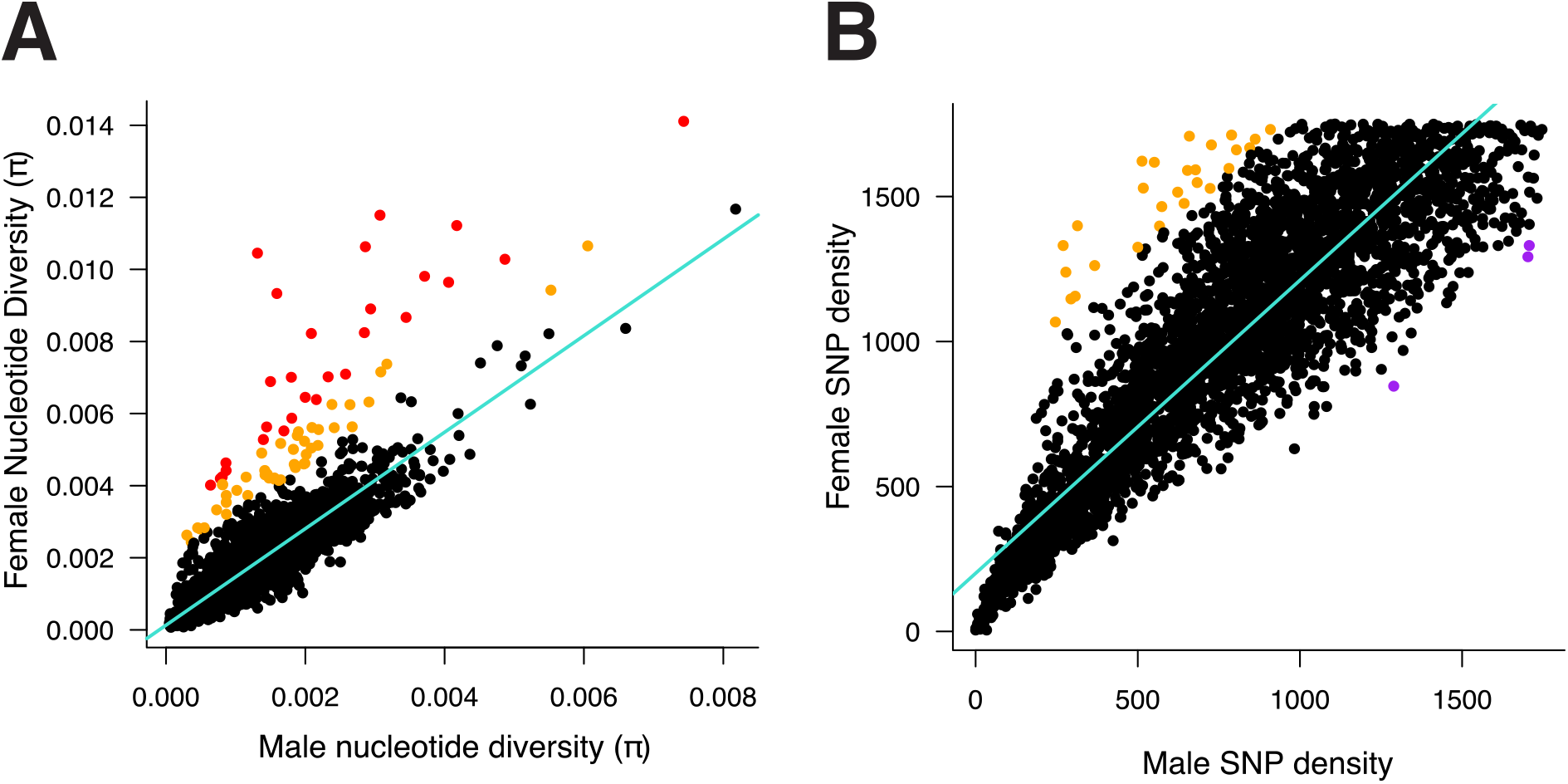
Comparison of genetic variation between females and males. (A) Nucleotide diversity (π) and (B) SNP density for males and females. Each point show the mean value of π or SNP density within a 50,000 bp window. A z-score was calculated using the distance from each point to the regression line (shown in teal). Points with z-scores > 3 are highlighted in orange, z-scores > 5 in red, and points with z-scores < -3 are highlighted in purple

Variation in genetic diversity could conceivably be driven by sampling differences among male and female samples rather than selective differences between the sexes. To test for the effect of sampling error on estimates of π, we generated 100 random subsamples of females and males (n=100 chromosomes/sex) and calculated the mean π and mean SNP density of each replicate (Figure S1). Consistent with the results from the full dataset, across all permutations, females had larger values of π (Welch’s t-test, t (163) = 50, P < 2.2e^-16^) and an increased number of SNPs (Welch’s t-test, t (197) = 60, P < 2.2e^-16^), indicating that sampling error had little effect on these results.

### Males have fewer deleterious variants than females

We investigated probable effects of SNPs and insertions/deletions (INDELs) variants in our dataset using the SnpEff (v4.3) (24) variant annotation and effect prediction tool. Females had more SNPs and INDELs than males both genome-wide and when considering only genic regions (Table 1). SNPs in genic regions were categorized as missense, nonsense, or silent variants. INDELs were classified by SnpEff as likely to have a high, moderate, low, or modifier effect on protein translation. Matching our predictions, males had a smaller proportion of missense and nonsense SNPs than females (two sample z-test: X^2^ = 7558, df = 2, P < 2.2e^-16^) as well as a lower proportion of high impact INDELs (two sample z-test: X^2^ = 1497, df = 3, P < 2.2e^-16^). Males had a higher transition/transversion (Ts/Tv) ratio than females (Table 1). Transversions are more likely to be detrimental than transitions (25), therefore this difference in Ts/Tv ratio could represent increased selection against transversions in the male population.

**Table 1:** SnpEff results for males and females

|  | FEMALES |  |  | MALES |
| --- | --- | --- | --- | --- |
| SNPS | Total Variants | 4,784,723 | Total Variants | 3,711,776 |
|  | Genic Variants | 302,883 | Genic Variants | 266,272 |
|  | Missense Variants | 117,439 (38.77%) | Missense Variants | 96,000 (36.05%) |
|  | Nonsense Variants | 3,294 (1.09%) | Nonsense Variants | 2,537 (0.95%) |
|  | Silent Variants | 182,150 (60.14%) | Silent Variants | 167,735 (62.99%) |
|  | Missense/Silent Ratio | 0.6447 | Missense/Silent Ratio | 0.5723 |
|  | Ts/Tv Ratio | 1.9015 | Ts/Tv Ratio | 2.0637 |
| IN/DELS | Total Variants | 1,743,723 | Total Variants | 1,344,848 |
|  | Insertions | 869,193 | Insertions | 685,520 |
|  | Deletions | 874,530 | Deletions | 659,328 |
|  | High Impact | 7,104 (0.29%) | High Impact | 4,664 (0.25%) |
|  | Moderate Impact | 4,776 (0.19%) | Moderate Impact | 4,002 (0.22%) |
|  | Low Impact | 5,129 (0.21%) | Low Impact | 2,822 (1.5%) |

To test for the effect of sampling error, we applied a similar permutation approach as above, repeating the SnpEff analysis for 100 random subsamples of 50 females and 100 males. The subsampled replicates produced similar results as the full dataset with males having a lower missense/silent ratio (Welch’s t-test, t (188) = 171, P < 2.2e^-16^), fewer nonsense variants (Welch’s t-test, t (116) = 59, P < 2.2e^-16^), and a greater Ts/Tv ratio than females (Welch’s t-test, t (104) = -128, P < 2.2e^-16^) (Figure S2).

### Males have fewer deleterious variants in housekeeping genes

If recessive deleterious alleles are preferentially exposed to selection in males, genes with deleterious variants in females but not in males may be enriched for genes necessary for development or survival. We investigated the identity of genes with nonsense variants in one sex but not in the opposite sex. We identified 773 genes with a nonsense variant in one or more of the female samples but without a nonsense variant in males (i.e., conserved in males). In the opposite direction, we identified 317 genes with a nonsense mutation in males but not females (i.e., conserved in females). Genes conserved in males were enriched for many gene ontology (GO) terms associated with basic organismal processes necessary for development such as mitotic spindle assembly checkpoint (GO:0007094), dorsal closure (GO:0007391), regulation of cell fate specification (GO:0042659), and mushroom body development (GO:0016319) (Table S2). Genes conserved in females were enriched for a wide range of GO terms including regulation of female receptivity (GO:0045924) (Table S3).

### Males have fewer low frequency variants than females

A third prediction of our hypothesis was that because most deleterious variants are likely at low frequencies, increased selection against deleterious variants in males is expected to lead to a reduction in the proportion of rare variants in males compared to females. To test this prediction, we first calculated the number of single copy variants (singletons) in our male and female datasets. The female dataset had 1,529,219 singletons whereas the male dataset had 1,309,484 singletons. Similar results were observed when comparing subsets of males and females with an equal number of chromosomes (Figure S3).

We next compared the distribution of rare nonsense and missense alleles between the sexes (Figure 3). For doubleton alleles (those with two copies in one of the sexes), we found that female doubletons were more likely to be missing in males than male doubletons were missing in females, illustrated by larger value of the red bar at a minor allele count of zero. Similar results were seen for alleles with 3-5 copies. This finding suggests that males lack many of the rare deleterious alleles that are in the female dataset. Synonymous variants in females showed a somwhat similar pattern as deleterious variants (Figure S4), which could result from background selection against deleterious variants removing linked synonymous variants.

**Figure 3:**
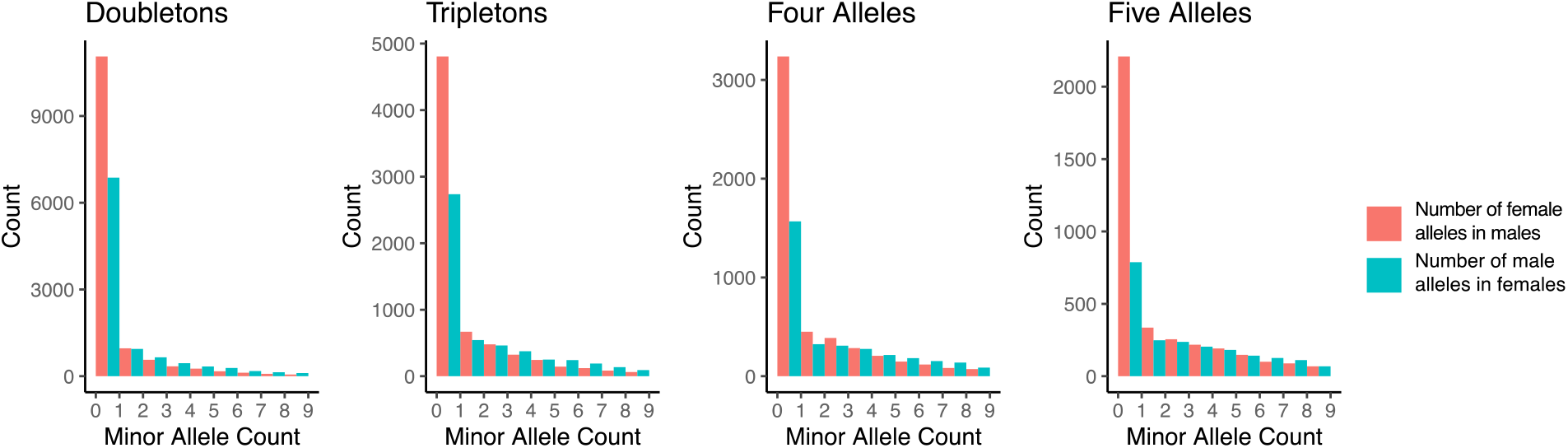
Comparison of rare nonsense and missense variants in one sex with the minor allele count in the opposite sex. Red shows the distribution minor allele count of female alleles in the male dataset. Blue shows the distribution minor allele count of male alleles in the female dataset. The red bar is greater than in blue bar for a minor allele count of zero indicating that variants with 2-5 copies in females are more often missing in males than variants with 2-5 copies in males are missing in females.

Finally, considering the distribution of nonsense, missense, and synonymous variants across all allele frequencies, we found that synonymous variants had a greater correlation between male and female allele counts (Slope = 0.88, R^2^ = 0.95), than missense (Slope = 0.85, R^2^ = 0.91) or nonsense variants (Slope = 0.70, R^2^ = 0.75) (Figure S5). This weaker correlation between allele count at nonsense variants is predicted if there is stronger selection against these deleterious alleles in males than females.

### Genetic load is lower in males

Our previous analyses considered allele frequency differences between males and females at the population level, but how does greater selection against deleterious alleles in the male population affect the genetic load of individuals? We estimated genetic load by calculating the number of nonsense and missense variants per individual. The average female is expected to have more variants than the average male due to the extra copy of each chromosome, therefore we compared the number of deleterious variants to the number of synonymous variants in each sample (Figure 4). The average male had 6.7% fewer synonymous variants than the average female. However, this reduction was even more pronounced for deleterious variants, with 17.7% fewer missense and 24.1% fewer nonsense variants in the average male than the average female. Male samples had a lower genetic load than female samples, consistent with our predictions.

**Figure 4:**
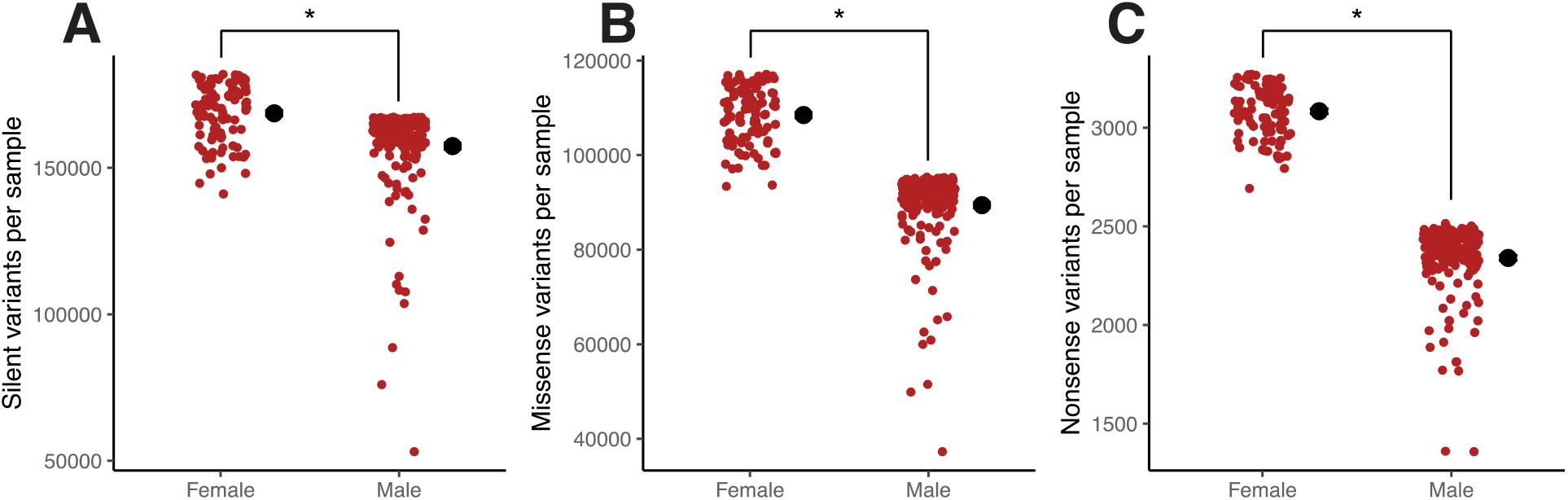
Genetic load lower in males. Using SnpEff, we calculated the number of (A) silent variants, (B) Missense variants, and (C) nonsense variants for each individual in our dataset.

## Discussion

In haplodiploid species, selection against deleterious alleles has been predicted to occur more frequently in haploid males than diploid females because recessive deleterious alleles in females can be masked from selection by partial or complete dominance. Consistent with this theoretical prediction, we found that haploid males have fewer overall variants, fewer deleterious variants, and fewer low frequency deleterious variants than diploid females. There are challenges to directly comparing allele frequencies between haploid and diploid groups due to differences in calling alleles for datasets with different ploidy, but any such bias would not explain the reduction in low frequency deleterious variants in males relative to females.

We considered several evolutionary mechanisms that might generate allele frequency differences between males and females that might explain our results. One possible mechanism might be a difference between male and female dispersal. A longer dispersal distance in one sex could increase the number of rare alleles detected in that sex. Prior comparisons of genetic diversity between the nuclear and mitochondrial genome suggest that males may be dispersing somewhat further than reproductive females (22). However, populations of *P. fuscatus* in Ithaca are panmictic (22) therefore gene flow from nearby locations will likely have little effect on allele frequency differences between the sexes. Moreover, longer-distance dispersal in males should increase the number of variants in males – the opposite effect of what we observed in our study – making dispersal differences an unlikely explanation for our results.

Differences in allele frequencies could reflect differences in effective population sizes between males and females. Although reproductive male and female *P. fuscatus* have a similar total number of individuals (19), females have a larger effective population size than males because females have twice the number of chromosomes. Natural selection is more efficient in larger populations, which could potentially lead to more efficient purging of deleterious mutations in females. However, more effective natural selection in females would lead to the inverse of our results, fewer genetic variants, and fewer deleterious variants, in females compared to males. Effective population size differences can also affect the number of new mutations contributed by each sex. With an equal mutation rate, new mutations will arise more often in diploid females than haploid males. Biases in certain nucleotide substitutions have been detected between haploid and diploid yeast (26) and an elevated mutation rate in females could also increase the number of rare alleles in females. However, in either case, the effect of increased mutational input from females will be short-lived. All mutations originating in females will be expressed in the next generation of males. The precise dynamics of the spread of new mutations in *P. fuscatus* depend on the mutation-selection balance in this species, which will require future study, but it seems unlikely that mutation rate differences alone are the primary cause of our results.

Another source of allele frequency variation between the sexes is sexually antagonistic selection (SAS), in which an allele is beneficial to one sex and harmful to the other sex (27, 28). It has been theorized that haplodiploids might be predisposed to extensive SAS and intralocus sexual conflicts in haplodiploids are predicted to be resolved in favor of females (29). In the tawny crazy ant (*Nylanderia fulva*), SAS for both male and female biased alleles was detected for ∼3% of the genome (30). However, SAS has not been extensively investigated in haplodiploids and it is unknown how widespread SAS is across haplodiploid species (27, 28). While SAS may explain some of the allele frequency differences between the sexes, it does not explain why allele frequency differences between the sexes are greatest for low frequency nonsense and missense variants.

The most likely and parsimonious explanation for our results is stronger purifying selection against deleterious mutations in haploid males. The magnitude of allele frequency differences between males and females at some sites in our dataset was quite large. In the most extreme case of selection, recessive lethal mutations would cause male mortality during development (Figure 1). Developing and dead larvae have been observed to be regularly ejected from nests (31), in one reported case, 58% of larvae were removed from a *P. dominula* nest (Pardi 1951 cited in 31). Researchers have not investigated the sex of removed larvae, but intriguingly larvae were rejected more frequently in the fall, which is when males are produced (31). Additionally, neutral, or deleterious variants linked to developmentally lethal alleles will also be removed from the male gene pool via background selection further decreasing genetic variation in the male population. Consequently, sisters may carry haplotypes, or even large chromosomal regions that are absent in their brothers.

Moderately or weakly deleterious alleles can impact male survival at later life stages, thereby further removing additional genetic variation from the sampled male population. Male *P. fuscatus* gather at leks or defend small territories near females (32), and many of our male specimens were collected at lekking sites. Males with a greater genetic load may be of poorer quality and therefore less capable of survival and competition, leading to earlier mortality, fewer mating opportunities, and stronger selection against males with deleterious alleles.

Haplodiploidy is a widespread method of sex determination, occurring in nearly 15% of arthropod species, with at least 14 independent origins of arrhenotokous haplodiploidy in insects (33, 34). If purifying selection is indeed stronger against haploid males than diploid females, this has several interesting implications for adaptation in haplodiploids. For example, the ant species *Formica aquilonia* and *Formica polyctena* hybridize in nature producing viable female offspring and inviable male offspring (36, 37), likely as a result of incompatibilities between recessive alleles in the two parental species. Selection against deleterious recessive alleles in haploid males is predicted to lower the genetic load and reduce inbreeding depression in haplodiploid species compared to diploid species (16, 17). This reduced genetic load may facilitate the development of mating systems with high levels of inbreeding such as the bark beetles (35). However, lower levels of genetic diversity in haplodiploid species could potentially slow rates of molecular evolution in comparison to diploid species. Alternatively, haplodiploids could compensate for lower genetic diversity through alternative mechanisms such as increased dispersal, or higher mutation or recombination rates. Indeed, comparisons among a limited number of species have found higher recombination rates in haplodiploid species compared to other insect species (38). Within haplodiploid species, haploid males may be more sensitive than females to the effects of environmental change, pesticide use, or disease. Future studies of environmental perturbations on haplodiploid insects may want to consider the effects separately between males and females. Comparisons between genetic variants present in diploid females but missing in haploid males has potential for rapidly identifying putative deleterious variants in natural populations.

## Materials and Methods

### Sample collection and sequencing

Male and female *P. fuscatus* were collected on the wing, at lekking sites, and on nests from sites around Ithaca, NY from 2015-2019 (Table S1). There is a high degree of gene flow across this region, with *P. fuscatus* forming a single panmictic population (22). Our dataset included all unrelated males collected (N=185). Female samples (N=100) were selected from a larger dataset of sequenced females from the Ithaca region. We chose female samples collected from similar collection sites as the male dataset, excluding females with kinship coefficients >0.1 as identified with the -relatedness2 option in VCFtools (v0.1.15) (39), and retaining samples with the highest depth of sequencing coverage. In some cases, males and females collected from the same nest were included in the dataset (Table S1).

DNA was extracted from wasp legs with the Qiagen Puregene Core Kit A. Individually barcoded paired-end whole-genome libraries were constructed with the Nextera library preparation kit with an average insert size of 550 bp. Library were sequenced on the Illumina HiSeq 2000 and NovaSeq 6000.

### Variant identification and filtering

Multiple filters were applied to sequenced data to remove variants caused by sequencing error rather than true biological variation. Raw sequencing reads were processed with Trimmomatic (v0.36) to remove adaptors and poor-quality sequence using the options SLIDINGWINDOW:4:15, LEADING:3, TRAILING:3, and MINLEN:36. Trimmed reads were aligned to the *P. fuscatus* reference genome (23) using the Burrows–Wheeler Aligner (v0.7.13) (40). Due to downstream software issues with having haploid and diploid variants in the same VCF file, variants were called separately in males and females. Variants were identified using Picard tools (v2.8.2; http://broadinstitute.github.io/picard) and the HaplotypeCaller tool in GATK (v3.8) (41). Variant calls for males and females were merged separately with the GenotypeGVCF tool in GATK, which aggregates variants across samples to correct genotype likelihoods and improve the confidence of variant identification. After alignment, poor confidence variants were hard filtered following GATK best practices recommendations. Single nucleotide polymorphisms (SNPs) were filtered using the parameters QUAL < 30, SOR > 3.0, FS > 60, MQ < 40, MQRankSum < -12.5, and ReadPosRankSum < -8.0. Insertion/Deletion (INDEL) variants were filtered using the parameters QD < 2.0, QUAL < 30, FS > 200, and ReadPosRankSum < -20. To remove additional variants caused by non-biological error or sites with poor coverage across the population, we produced a final filtered male and female dataset by removing variants with more than 80% missing data, and variants with exceptionally low or high depth of sequencing coverage across all individuals, using the filters --min-meanDP 3, --max-meanDP 500, and -- max-missing 0.2 in VCFtools (39).

### Measurement of genetic variation across the genome

Using the filtered male and female SNP datasets, we calculated the mean nucleotide diversity (π) and mean SNP density in 50,000 bp window using VCFtools (39). We plotted the mean value of π or SNP density for each sex, and then calculated a z-score for each window using the distance from each point to the regression line. Windows with z-scores > 3 or < -3 were considered outlier windows. Postitive z-scores indicate that π or SNP density was higher in females then expected.

We used a permutation approach to test for the possible effect of sampling error on π and SNP density. In brief, we randomly subsampled 50 females and 100 males from each dataset (equal number of chromosomes between the sexes) and calculated the mean genome-wide π and SNP density. We repeated this for 100 random subsets of males and females. Differences between the male and female subsamples were assessed using t-tests implemented in R (v4.0.2) (42).

### Predicting variant effects and estimating genetic load

To predict the probable effects of variants in our dataset, we used the SnpEff (v4.3) (24) variant annotation and functional effect prediction tools on the filtered female and male SNP and INDEL datasets. To assess the possible effect of sampling error on these results, we repeated SnpEff on the same 100 subsets of 100 females or 50 males in the permutation test described above.

Using the output of SnpEff, we assembled a list of genes containing at least one nonsense variant in one or more of the females in our dataset but lacking nonsense variants in the male dataset (“conserved in males”). For genes conserved in males, we looked for enrichment of Gene Ontology (GO) terms using the TopGO package in R (43), choosing the ‘weight01’ algorithm to incorporate the GO hierarchy when identifying enriched GO terms, and assessing significance using fisher’s exact test. We repeated this analysis for genes without nonsense variants in females but with at least one nonsense variant in our male dataset (“conserved in females”).

To estimate genetic load of each sample, we used SnpEff to identify the number of missense, nonsense, and synonymous variants in each individual sample.

### Distribution of low frequency variants

We used VCFtools (39) to calculate the number of single copy alleles in the complete male and female datasets and in the 100 subsets of 100 females or 50 males. We then compared the distribution of nonsense, missense, and synonymous alleles between the male and female datasets. We compared the number of alleles rather than the allele frequency to avoid inflating allele frequencies for sites with lower coverage across samples. Variants were filtered by effect using SnpSift (v4.3) (24) and the minor allele count was calculated at each site using VCFtools (39). As the original VCF file contained only variable sites, variants that were missing in the opposite sex could be caused by lack of sequence data at that site, or because that site is invariant in the opposite sex. To distinguish between these possibilities, we output all invariant sites in the male and female dataset using GATK and kept only sites that were present in both sexes.

