## Supplemental Figures for "Sex differences in deleterious genetic variants in a haplodiploid social insect"

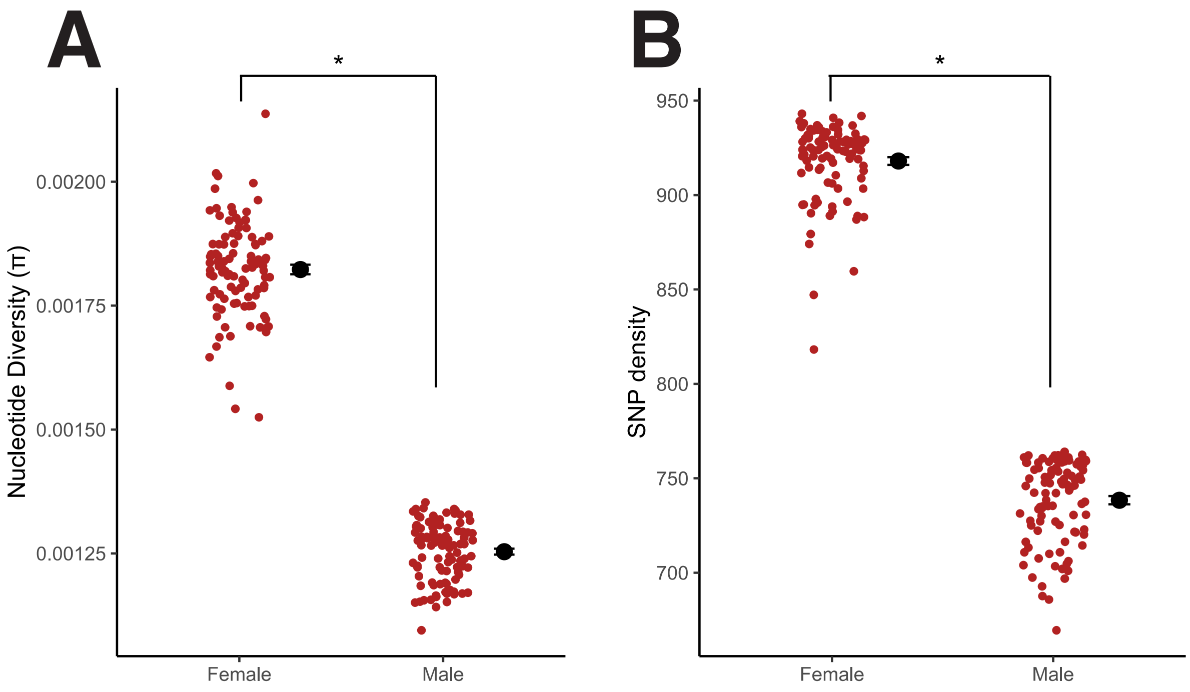


Figure S1: Variation in nucleotide diversity (π) and SNP density between sexes not due to sampling error. We used a permutation approach to test for the effect of sampling error on estimates of genetic diversity by subsampling 50 random females and 100 random males (n=100 chromosomes/sex). Each point shows the (A) mean genome-wide π or (B) mean SNP density calculated for 100 subsamples. Females had significantly larger π (Welch's t-test, $t\ (163) = 50$, $P < 2.2e^{-16}$) and a higher density of SNPs (Welch's t-test, $t\ (197) = 60$, $P < 2.2e^{-16}$) for all permutations. The mean and standard error of each estimate are shown to the right of each set of permutations.

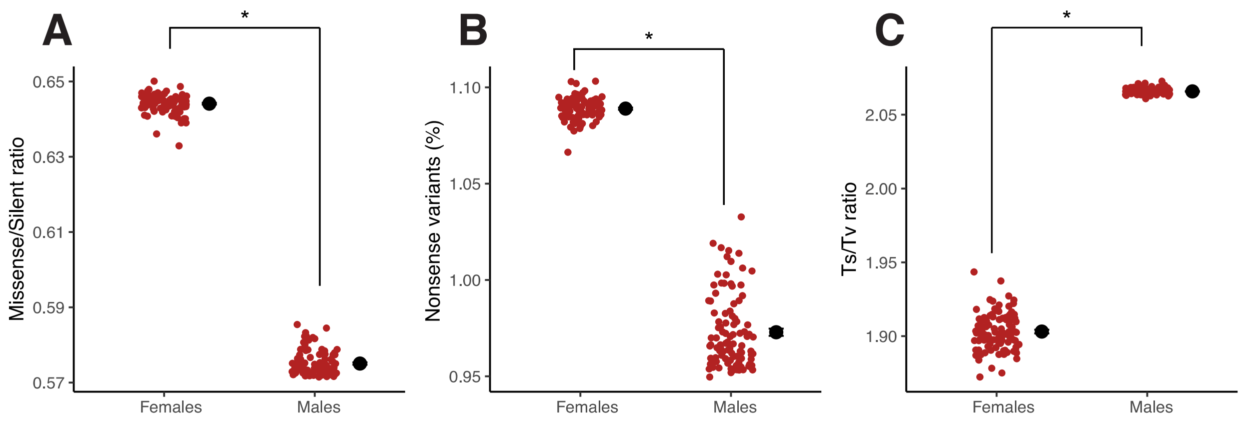


Figure S2: SnpEff permutation results. To test for the effect of sampling error, we subsampled 50 random females and 100 random males (n=100 chromosomes/sex) for 100 permutations. Each point shows a single replicate, and the mean and standard error of each sex are shown to the right of each set of permutations. Females had (A) a higher ratio of missense to silent variants (Welch's t-test, t (188) = 171, P < $2.2e^{-16}$), (B) a greater proportion of nonsense mutations (Welch's t-test, t (116) = 59, P < $2.2e^{-16}$), and (C) a lower transition/transversion ratio (Ts/Tv) (Welch's t-test, t (104) = -128, P < $2.2e^{-16}$).

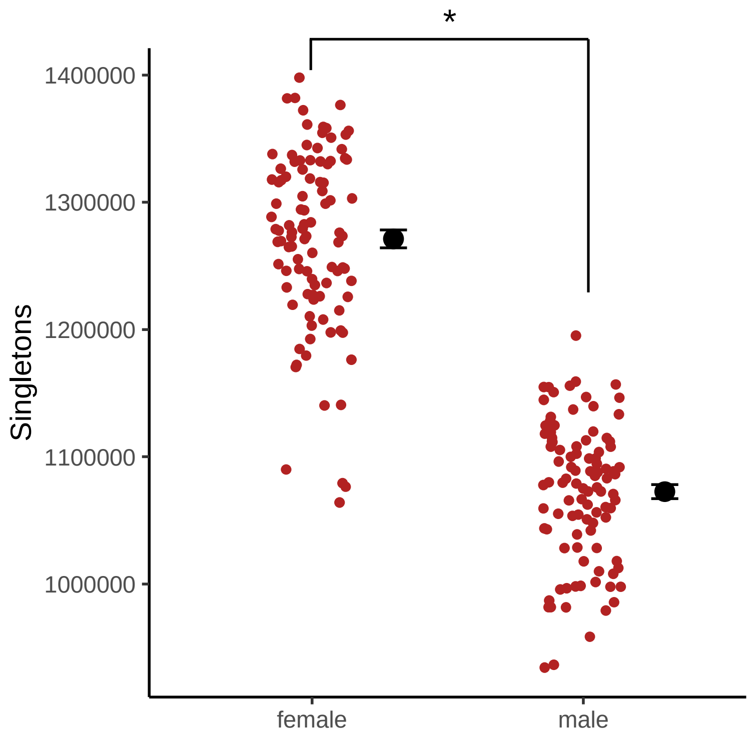


Figure S3: Males have fewer singletons than females. We subsampled 50 random females and 100 random males (n=100 chromosomes/sex) for 100 permutations. For each permutation, we calculated the number of single copy variants (singletons) in each sex. Points show a single replicate. The mean and standard error of each sex are shown to the right of each set of permutations.

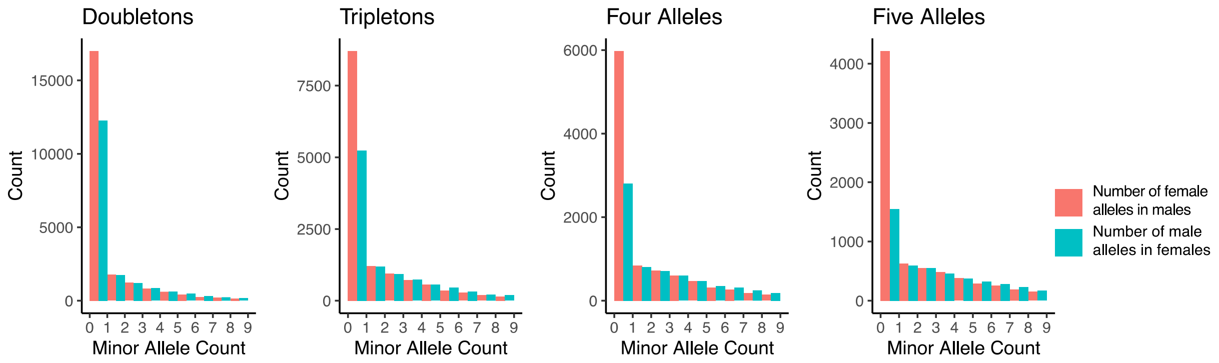


Figure S4: Comparison of rare synonymous variants in one sex with the minor allele count in the opposite sex. Red shows the distribution minor allele count of female alleles in the male dataset. Blue shows the distribution minor allele count of male alleles in the female dataset. The red bar is greater than in blue bar for a minor allele count of zero indicating that variants with 2-5 copies in females are more often missing in males than variants with 2-5 copies in males are missing in females.

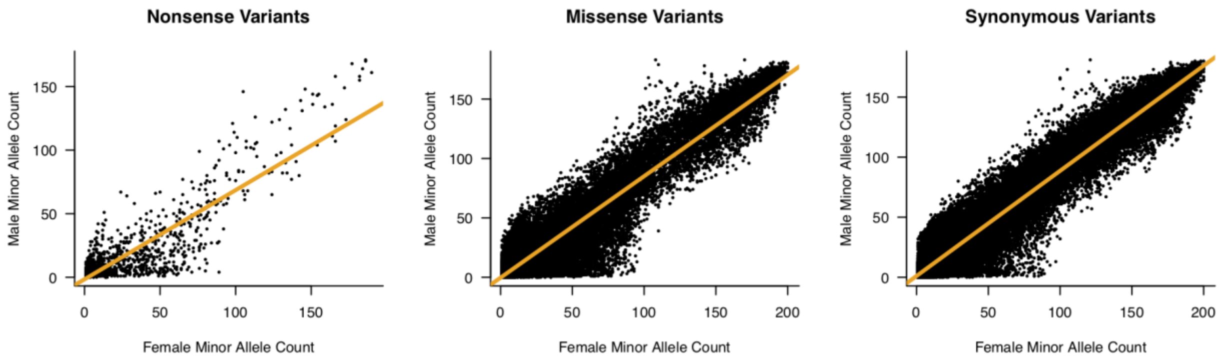


Figure S5: Male versus female minor allele count. Each point shows the minor allele count in each sex for a single SNP. The correlation between male and female minor allele count for (A) nonsense variations (Slope=0.70, $R^2$= 0.752) is weaker than for (B) missense variants (Slope=0.85, $R^2$= 0.907) or (C) synonymous variants (Slope=0.88, $R^2$ = 0.945). The line is the regression line for the two values.
